# Learning-dependent cholinergic plasticity reconfigures cortical circuit dynamics

**DOI:** 10.1101/2025.11.10.687664

**Authors:** Andrew H. Moberly, Jessica A. Cardin, Michael J. Higley

## Abstract

Neuromodulation by acetylcholine (ACh) plays a critical role in shaping neural dynamics in the neocortex as a function of development, behavioral state, and learning ^1–6^. Prior work suggests cholinergic signaling can act as a gate for the subsequent induction of circuit plasticity ^3,7,8^. However, modification of ACh release could also be a direct mechanism for the expression of cortical plasticity. Here, we combine widefield and 2-photon imaging in head-fixed mice to show that visual fear conditioning leads to a selective, cue-dependent release of ACh in primary visual cortex that enhances visually-evoked neuronal responses via excitation of layer 1 GABAergic interneurons and resulting disinhibition of local excitatory pyramidal neurons. Cholinergic signaling through muscarinic receptors in visual cortex is necessary for both learning and expression of conditioned fear behavior and enhanced visual representation. Our results demonstrate a novel capacity for conditioned release of ACh in sensory cortex to serve as a mechanism for sensory-guided behavioral learning. Rather than acting as a simple gate, cortical neuromodulation may thus play a central role in the expression of learned behavior.

## Main Text

Release of neuromodulators such as acetylcholine (ACh) is coupled to spontaneous variation in behavioral state and dynamically reconfigures local and long-range circuit organization across the neocortex^4,9–12^. Cholinergic release also occurs following delivery of rewarding and aversive stimuli, potentially playing a role in learning-dependent plasticity in cortical networks ^2,4,9,11,13–18^. A rich literature demonstrates that learning to associate sensory cues with reinforcement or punishment modifies neural representations of behavioral task variables and alters the sensitivity of responses to subsequent external inputs ^19–26^. This network-level plasticity may reflect the reorganization of local microcircuits, including subpopulations of GABAergic interneurons, many of which express cholinergic receptors ^27–29^. Indeed, cholinergic modulation of cortical inhibition is implicated in state-dependent cortical activity ^13,27,30,31^. However, the potential for direct learning-related plasticity of cortical ACh release in sensory areas and the consequences for circuit dynamics and behavior remain unexplored.

Classical fear conditioning involves repeated pairing of a neutral sensory cue with an aversive stimulus such as a mild foot-shock ^32,33^. In primary auditory cortex, fear conditioning can enhance sensory-evoked activity of layer 2/3 excitatory pyramidal neurons (PNs) and shift their tuning properties towards the conditioned frequency ^34–36^, a form of plasticity that requires activation of muscarinic ACh receptors ^5,6,8,37,38^. Indeed, foot-shock drives acute ACh release in auditory cortex of untrained anesthetized animals, potentially disinhibiting PNs via activation of layer 1 GABAergic interneurons (L1-INs) that in turn suppress layer 2/3 parvalbumin-expressing interneurons (PV-INs) ^3^. This acute disinhibition is thought to gate the induction of learning-dependent cortical plasticity ^39–42^. However, the dynamics and plasticity of ACh release itself during task learning and their role in the acquisition and expression of learned behavior are unknown.

To investigate how learning modifies both ACh release and the functional organization of cortical microcircuits, we developed a novel visually-cued fear conditioning task in which head-fixed mice learn to associate a brief visual stimulus with a mild foot-shock (Figure 1a). Animals were first water-deprived, acclimated to head-fixation, and trained to lick a spout for freely available water. We then presented a visual stimulus comprising a 100% contrast, 5s full-screen, dynamic filtered noise pattern for 10-20 trials per day with a random intertrial interval (ITI, Extended Data Figure 1, see Methods). Animals were shown the visual stimulus alone for three days to assess baseline licking behavior. Beginning on the fourth day, the visual stimulus was followed after a 1s gap by a mild foot-shock that resulted in immediate cessation of licking behavior. Repeated stimulus-shock pairings rapidly induced learned lick suppression at the onset of the visual cue that lasted for the duration of the stimulus presentation and persisted for several seconds (Extended Data Figure 1). We quantified licking behavior using a lick rate index calculated over the 5s visual stimulus duration (Rate_Post_-Rate_Pre_)/(Rate_Post_+Rate_Pre_). This value decreased immediately on the first day of conditioning and reached a plateau within 2-3 days (Figure 1b), yielding an overall significant reduction in lick index for the population (Pre vs. Post lick index, −0.11±0.04 vs. −0.76±0.12, p<0.01 paired t-test, n=17 mice, Figure 1c). Learned lick suppression was robust for high contrast stimuli but decreased with reduced visual contrast (Extended Data Figure 1), consistent with expected variation in perceptual detectability. In addition to lick suppression, conditioning also produced a significant increase in pupil dilation associated with the onset of the visual stimulus (Pre vs. Post pupil area, −0.02±0.13 ΔZ vs. 0.64±0.13 ΔZ, p<0.01 paired t-test, Figure 1f-i). Head-fixed mice thus readily learn an association between a previously neutral visual cue and an aversive unconditioned stimulus, resulting in both somatic and autonomic conditioned responses.

**Figure 1:**
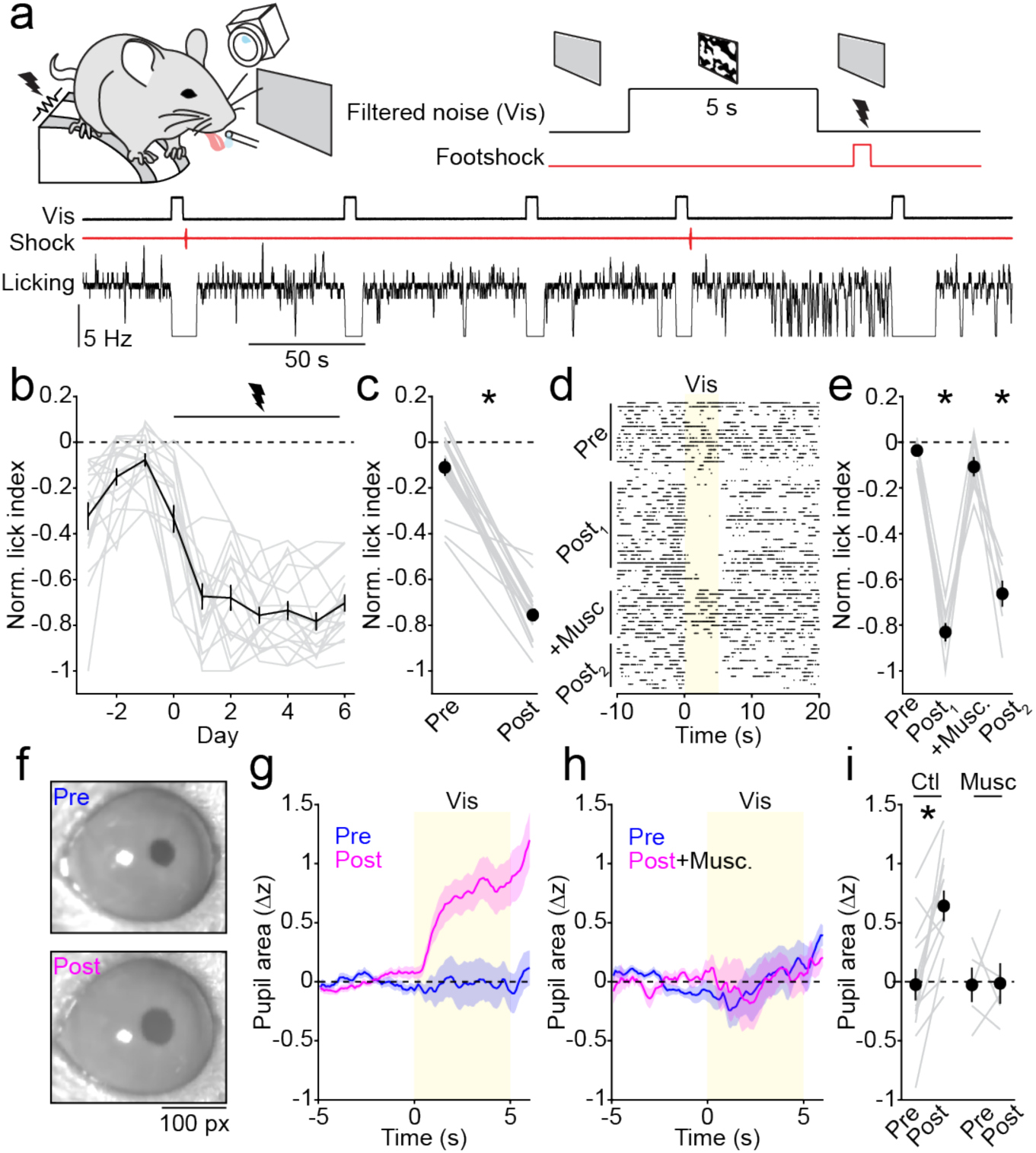
Visually cued lick suppression is rapidly induced and dependent on visual cortex activity. (**a**) Schematic of the behavioral setup. During conditioning, a 5-second filtered noise stimulus is followed by a 1-second trace and a 0.5-second foot shock. Example traces show visual stimulus presentation, foot shock, and lick rate from a portion of a post-conditioning session. (**b**) Normalized lick indices per day, with day 0 marking the first pairing of visual stimulus and foot shock (n=17 mice). (**c**) Average pre-and post-conditioning lick indices (n=17 mice; individual mouse averages in gray) * indicates p<0.05, paired t-test. (**d**) Example lick raster aligned to visual stimulus onset across pre-(Pre) and post-conditioning (Post_1_), muscimol session (+Musc), and post-muscimol session (Post_2_). (**e**) Average pre, post conditioning, muscimol session, and post muscimol session conditioning lick indices (n=7 mice) * indicates p<0.05 for pairwise post-hoc comparisons (Tukey-Kramer corrected) against pre-conditioning session following a significant (p<0.05) main effect of session in repeated measures ANOVA. (**f**) Example pupil images representing the average frame across the 5-second visual stimulus from a pre-(Upper) and post-(Lower) conditioning session. (**g**) Averaged pupil traces (n=13 mice) aligned to the visual stimulus from pre-(blue) and post-(magenta) conditioning sessions. (**h**) Averaged pupil traces (n=13 mice) aligned to the visual stimulus from pre-and post-conditioning muscimol sessions. (**i**) Pre- and post-conditioning pupil area from sessions without muscimol (Ctl, n=13 mice) and muscimol sessions (Musc, n=5 mice). * indicates p<0.05, paired t-test. Data in panels **b**, **c**, **e**, **g**, h,& **i** are represented as mean ± SEM.

Conditioning did not significantly change the baseline lick rate during the ITI nor the basal arousal level associated with head-fixation or placement in the experimental setup, as indicated by the lack of difference in the average ITI pupil size before and after stimulus-shock pairing (Extended Data Figure 1). Similar levels of learned lick suppression were observed across multiple independent cohorts of mice used throughout the present study (Extended Data Figure 1), demonstrating the reproducibility of this assay for quantifying visually cued conditioning.

Normal activity in primary visual cortex (VisP) was necessary for the expression of learning, as acute intracortical injection of the GABA receptor agonist muscimol reversibly abolished conditioned lick suppression (F=88.92, p<0.01, ANOVA; p<0.01 Tukey’s Post-hoc for muscimol vs. control, n=7 mice, Figure 1d-e) with no change in basal lick rate during the ITI (Extended Data Figure 1). Moreover, inactivation of VisP also abolished the visually-evoked pupil dilation after learning (Pre vs. Post pupil area, −0.03±0.15 ΔZ vs. −0.01±0.17 ΔZ, Figure 1h-i). Overall, these results demonstrate that the neocortex is a critical mediator of visually cued somatic and autonomic behavior.

To explore how this form of learning impacts the cortical representation of the visual cue, we assessed cortical activity using widefield, mesoscopic calcium imaging ^4,43,44^. We used neonatal injections of adeno-associated virus (AAV) into the cerebral vasculature to induce cortex-wide expression of the green fluorescent calcium indicator GCaMP6s ^4,45–47^. Adult mice were then trained as described above while under the microscope objective (Figure 2a). Mesoscopic imaging through the intact skull generated robust signals across the dorsal neocortex, sufficient for monitoring both spontaneous and visually evoked activity (Figure 2b-c). We used a series of drifting filtered-noise bars (see Methods) to map the retinotopic borders of VisP and higher-order visual areas based on fluorescent calcium signaling (Figure 2b), allowing us to quantify neural activity from identified regions during task performance. Individual trials showed a characteristic visually-evoked response that began in VisP and rapidly spread throughout much of the cortex, as in previous work ^48^ (Figure 2d-e).

**Figure 2:**
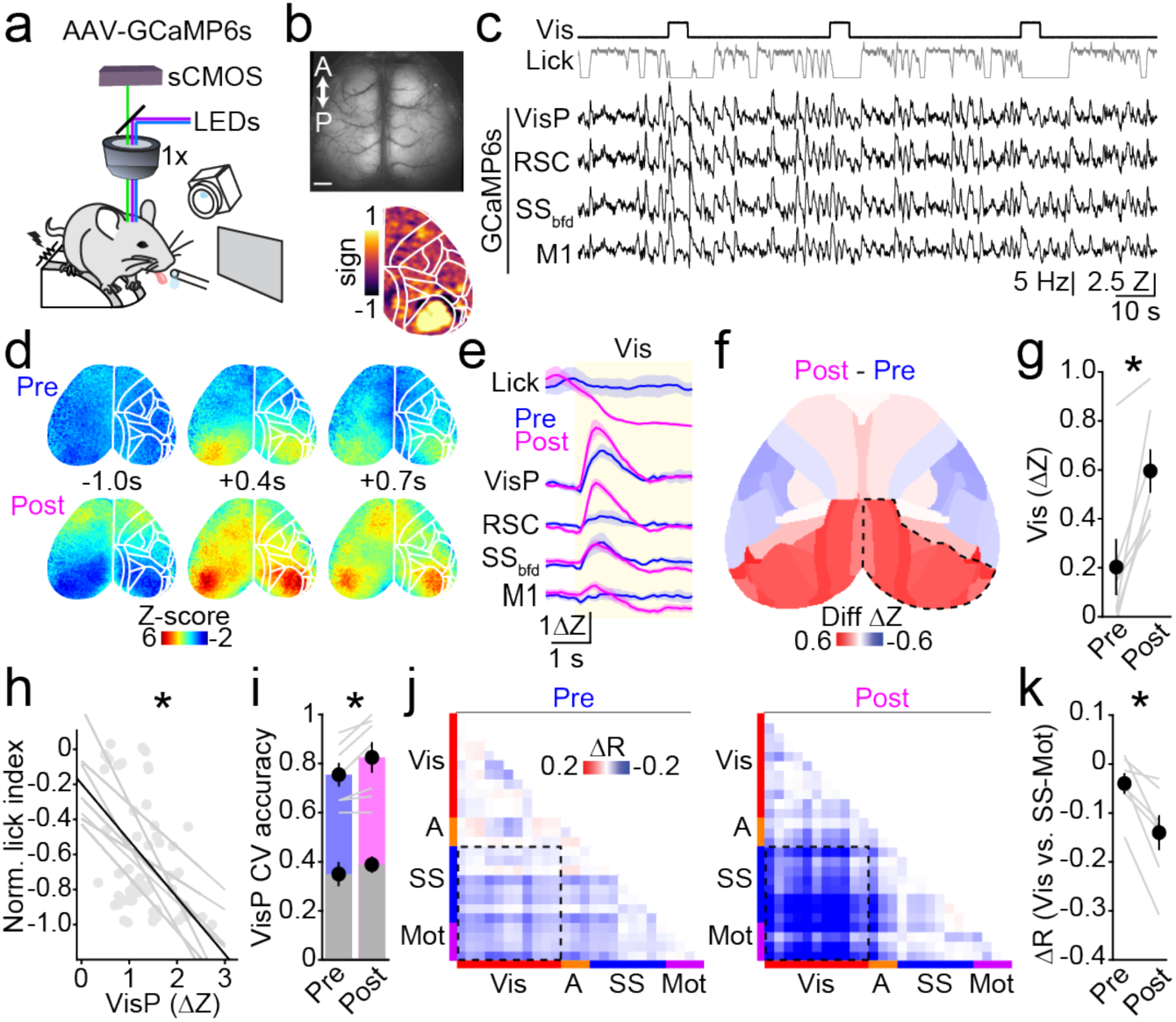
Visual fear-conditioning enhances visually-evoked activity. (**a**) Schematic of the widefield imaging setup. (**b**) Upper: example transcranial imaging field of view showing GCaMP6 expression. Scale bar: 1mm. Lower: averaged retinotopic field sign maps with Allen CCFv3 parcels overlaid in white. (**c**) Example time series from a portion of a post-conditioning session showing onsets of visual stimuli, lick rate, and traces corresponding to calcium activity in Allen parcels (VisP; primary visual cortex, RSC; retrosplenial cortex, SS_bfd_; somatosensory cortex barrel field, M1; primary motor cortex). (**d**) Example image frames showing a visual stimulus response pre- and post-conditioning from the same subject. White lines indicate Allen CCFv3-derived parcellation. Times in seconds are relative to visual stimulus onset at t=0s. (**e**) Traces show visual stimulus-averaged activity pre-(blue) and post-(magenta) conditioning from the Allen parcels in (**c**) (n=7 mice). (**f**) Average spatial map of the difference (post minus pre-conditioning) in the visual response amplitude for each Allen parcel (n=7 mice). Red indicates increased response amplitude and blue indicates decreased response amplitude. (**g**) Average response across visual parcels, represented by dashed outline in (**f**), pre- and post-conditioning (n=7 mice) * indicates p<0.05, paired t-test. (**h**) Relationship between normalized lick index and evoked visual response; points represent individual sessions with associated best-fit lines from each animal. The black line represents the average (n=7 mice). * indicates p<0.05, linear mixed-effects model. (**i**) 10-fold cross-validated SVM accuracy classifying pre-stimulus baseline vs. visually-evoked activity (VisP parcel). Gray bars: cross-validated SVM accuracy trained on time-shuffled data. * indicates p<0.05, paired t-test. (**j**) Pairwise correlation matrices between Allen Parcels representing the difference between values during visual stimuli and values outside of the visual stimulus (i.e., interstimulus intervals) pre-(left) and post-(right) conditioning. Matrices are averaged over 7 mice. Red indicates increased correlations and blue indicates decreased correlations. (**k**) Averaged correlation differences between visual and somatosensory/motor areas (dashed area outlined in **j**). * indicates p<0.05, paired t-test. Data in panels **e**, **f**, **g**, **i** & **k** are represented as mean ± SEM.

Imaged mice exhibited robust learning-induced lick suppression (Extended Data Figure 1). We compared visually-evoked signals before and after conditioning, finding a substantial increase in the magnitude of the visual response (Figure 2d-f, Extended Data Figure 2). The evoked response across visual areas increased significantly with learning (Pre vs. Post evoked response, 0.20±0.11 ΔZ vs. 0.60±0.09 ΔZ, p<0.05, paired t-test, n = 7 mice, Figure 2g, Extended Data Figure 2). However, there was no significant difference in the evoked response for somatosensory-motor regions (Extended Data Figure 2), suggesting selective plasticity in visual cortex. As a control experiment, we found that averaging cortical signals around spontaneous periods of lick cessation showed no increased activity in visual areas and revealed a modest reduction within frontal motor areas (Extended Data Figure 2), suggesting that the conditioning-induced increase in visual response was not related to altered motor behavior.

We next examined whether the increased visual response was directly associated with improved perceptual performance. First, we observed that across sessions, the average evoked response was significantly linearly related to the magnitude of lick suppression (β = −0.21, SE = 0.08, t(56) = −2.81, p = 0.007, linear mixed effects model, n=7 mice, Figure 2h). We also found that the accuracy of a linear decoder to detect a visual stimulus based on VisP activity significantly improved after conditioning (Pre vs. Post classification accuracy, 0.76±0.05 vs. 0.82±0.06, p<0.05, paired t-test, n=7 mice, Figure 2i), suggesting that the learning-induced plasticity may enhance detection of the cue.

Time-varying changes in behavioral performance were also associated with fluctuations in functional connectivity across large-scale cortical networks^4,43^. Indeed, prior to conditioning, presentation of the visual stimulus induced a modest decorrelation between visual and somatosensory-motor areas (Pre-inter-trial period minus visual stimulus ΔR, −0.04±0.02, Figure 2j). Learning induced a significant enhancement of this stimulus-induced decorrelation (Post-inter-trial period minus visual stimulus ΔR, −0.14±0.04, p<0.05, paired t-test, n=7 mice, Figure 2j-k), though we dd not observe a change in cortical functional connectivity during spontaneous activity in the inter-trial periods (Extended Data Figure 2). These results further indicate that learning induces selective plasticity in the cortical response to visual input.

To further test whether learning-related cortical plasticity depends on task contingencies, we examined whether this selectivity occurred for specific visual cues. In a separate cohort of animals, we found that mice trained on a task variant with distinct visual cues (CS+ and CS-, paired and unpaired with shock, respectively) demonstrated lick suppression only to the CS+ (Extended Data Figure 2). Moreover, training significantly enhanced the visual response to the CS+ relative to the CS-(Extended Data Figure 2). Thus, visually cued fear conditioning results in a selective enhancement of the cortical response to the conditioned stimulus.

Mesoscopic imaging data reflect spatially averaged signals arising from cell bodies and the surrounding neuropil ^45,49–52^. Therefore, to determine whether learning produces a specific increase in the activity of individual cortical neurons, we carried out 2-photon calcium imaging in a separate cohort of transgenic mice conditionally expressing GCaMP6s in excitatory VGlut1-expressing PNs ^53^ (Figure 3a). Mice were implanted with a titanium headpost and glass cranial window over VisP and behaviorally conditioned as described above. For each mouse, we initially used mesoscopic imaging to identify the borders of VisP, followed by 2-photon imaging of individual layer 2/3 PNs (L2/3 PNs) (Figure 3b). As above, training produced significant lick suppression across the group (Extended Data Figure 1). Consistent with our mesoscopic data, conditioning was associated with a significant overall increase in the magnitude of the visually-evoked response averaged across individual L2/3 PNs (Pre vs. Post evoked response, 0.51±0.13 Z vs. 0.82±0.22 Z, β = −0.19, SE = 0.03, t(28145) = −6.75, p<0.001, linear mixed effects model, n=8 mice, Figure 3c-e). Taken together, our mesoscopic and 2-photon imaging results indicate that visual fear-conditioning induces a significant, stimulus-specific enhancement of PN sensory representation in VisP.

**Figure 3:**
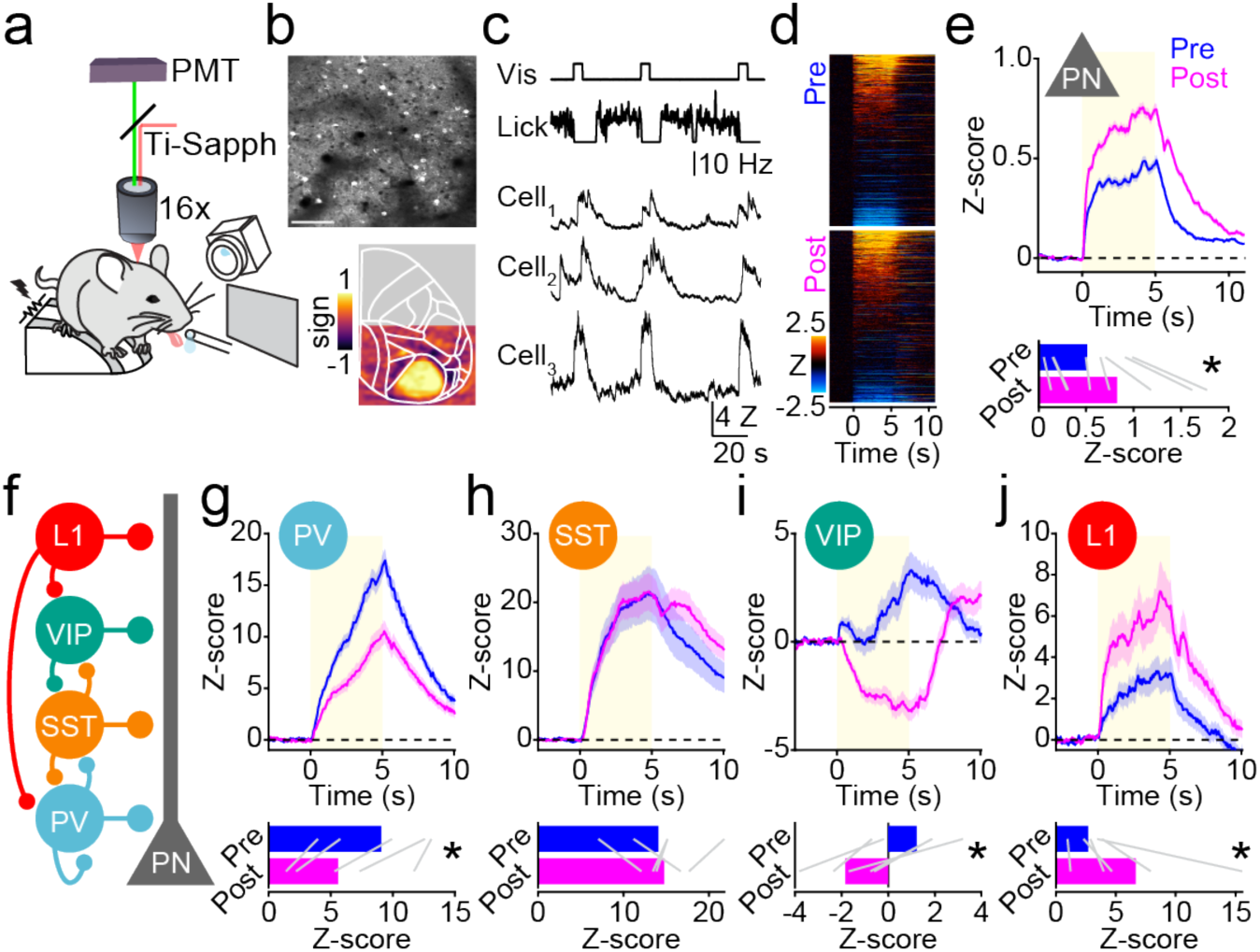
Plasticity of local circuit dynamics following visual fear conditioning. (**a**) Schematic of the 2-photon imaging setup. (**b**) Upper: example 2-photon imaging field of view. Scale bar, 100 μm. Lower: averaged retinotopic field sign maps with Allen CCFv3 parcels overlaid in white. (**c**) Example time series from one animal from a portion of a post-conditioning session showing onsets of visual stimuli, lick rate, and traces from three layer 2/3 PNs. (**d**) Evoked responses from all layer 2/3 PNs aligned to visual stimuli pre- and post-conditioning. (**e**) Upper: Averaged stimulus-aligned visual response traces pre- and post-conditioning from all excitatory cells. Lower: Average response amplitude pre-(blue) and post-(magenta) conditioning averaged across animals (n=8 mice). * indicates significant main effect of condition (pre vs post conditioning) on response amplitude in a linear mixed effects model. (**f**) Schematic of local cortical circuitry. (**g**-**j**) Same as (**e**) for PV-INs, SST-INs, VIP-INs, and L1-INs (n =5 PV-Cre mice; n=5 SST-Cre mice; n=5 VIP-Cre mice; n=5 mDLX-GCaMP6f L1 mice). * indicates significant main effect of condition (pre vs post conditioning) on response amplitude in a linear mixed effects model. Data in panels **e**, **g**, **h**, **i** & **j** are represented as mean ± SEM.

Visually-evoked activity in the neocortex is strongly shaped by the interplay between excitatory PNs and diverse subpopulations of GABAergic INs ^54^ (Figure 3f). In particular, PV-INs play a well-established role in constraining the magnitude and timing of PNs output ^55–58^. We tested the hypothesis that enhanced visual responses might be associated with a weakening of PV-IN activity following learning. We expressed GCaMP6f in PV-INs of VisP via local AAV injection (see Methods) and carried out 2-photon imaging of these cells during conditioning (Extended Data Figure 3). As above, these mice readily learned to associate the visual cue with shock (Extended Data Figure 1). However, in contrast to results from PN imaging, conditioning induced a significant reduction in the magnitude of visual responses in PV-INs (Pre vs. Post evoked response, 9.08±1.82 Z vs. 5.60±1.86 Z, β = 3.80, SE = 0.73, t(202) = 5.23, p<0.001, linear mixed effects model, n=5 mice, Figure 3g). In addition to PV-INs, GABAergic interneurons expressing either somatostatin-(SST-INs) or vasoactive intestinal peptide (VIP-INs) play key roles in shaping the activity of PNs in VisP ^30,59–63^ (Figure 3f). Therefore, we used a similar strategy to image the visually evoked responses in these cell types before and after conditioning. Mice expressing GCaMP6f in either SST-INs or VIP-INs reliably exhibited conditioned lick suppression (Extended Data Figure 1). While SST-INs did not show a significant change in visual response associated with training (Pre vs. Post evoked response, 14.08±2.25 Z vs. 14.76±1.05 Z, β = −0.12, SE = 3.13, t(162) = −0.04, p=0.97, linear mixed effects model, n=5 mice, Figure 3h), VIP-IN responses were significantly reduced below baseline levels (Pre vs. Post evoked response, 1.22±0.79 Z vs. −1.84±0.59 Z, β = 3.10, SE = 0.62, p<0.001, linear mixed effects model, n=5 mice, Figure 3i). Fear conditioning thus selectively induces a broad suppression of sensory-evoked GABAergic output in two major classes of VisP interneurons.

Recent studies have suggested that GABAergic interneurons residing in Layer 1 can provide blanket inhibition to multiple postsynaptic IN classes ^39,64–68^ (Figure 3f). We reasoned that these cells might be well-positioned to enhance visual responses in PNs via inhibition of local GABAergic microcircuits. To explore this hypothesis, we expressed GCaMP6s in VisP via AAV with an interneuron-specific enhancer ^69^ and carried out 2-photon imaging of cell bodies in layer 1 (Extended Data Figure 3). Again, these mice readily learned the task (Extended Data Figure 1). In contrast to the other interneuron subpopulations, conditioning produced a significant increase in the magnitude of the L1-IN visually evoked response (Pre vs. Post evoked response, 2.69±0.50 Z vs. 6.67±2.47 Z, β = −3.06, SE = 1.08, t(260) = −2.84, p=0.005, linear mixed effects model, n=5 mice, Figure 3j). Combined with our other imaging data, these results suggest that visual fear conditioning induces a significant enhancement of sensory-evoked activity in L1-INs, leading to a suppression of activity in both PV- and VIP-INs and a disinhibitory increase in PN output.

Previous work found that L1-INs are highly sensitive to cholinergic signaling ^3,31,70^, and moreover cholinergic modulation is strongly implicated in behavioral state- and learning-dependent variation in cortical circuit dynamics ^6,9,71–75^. We therefore hypothesized that plasticity of cortical ACh release might contribute to the functional reorganization of visual representation. To explore this possibility, we combined neonatal AAV injection to drive pan-cortical expression of the green fluorescent ACh sensor GRABS_ACh_ ^4,76^ with mesoscopic imaging to monitor cholinergic dynamics before and after conditioning (Figure 4a). Again, hemodynamic imaging allowed us to identify visual areas in these animals (Figure 4b). Imaging in untrained mice revealed spatiotemporally dynamic and heterogeneous spontaneous ACh release across the cortex and a weak signal in visual areas associated with stimulus onset (Figure 4c-e), consistent with prior reports ^77,78^. However, after training, presentation of the visual cue elicited robust ACh release (Figure 4e-f). As with the calcium response, learning induced a significant increase in the stimulus-evoked cholinergic signal in visual areas (Pre vs. Post evoked response, 0.11±0.07 ΔZ vs. 0.32±0.22 ΔZ, p<0.05, paired t-test, n=7 mice, Figure 4g), but not somatosensory-motor areas (Extended Data Figure 4). The increased ACh release was not due to reduced motor activity per se, as spontaneous lick pauses did not produce measurable cholinergic signals (Extended Data Figure 4). Visually cued fear learning is thus associated with a selective enhancement of cholinergic release in visual cortical areas.

**Figure 4:**
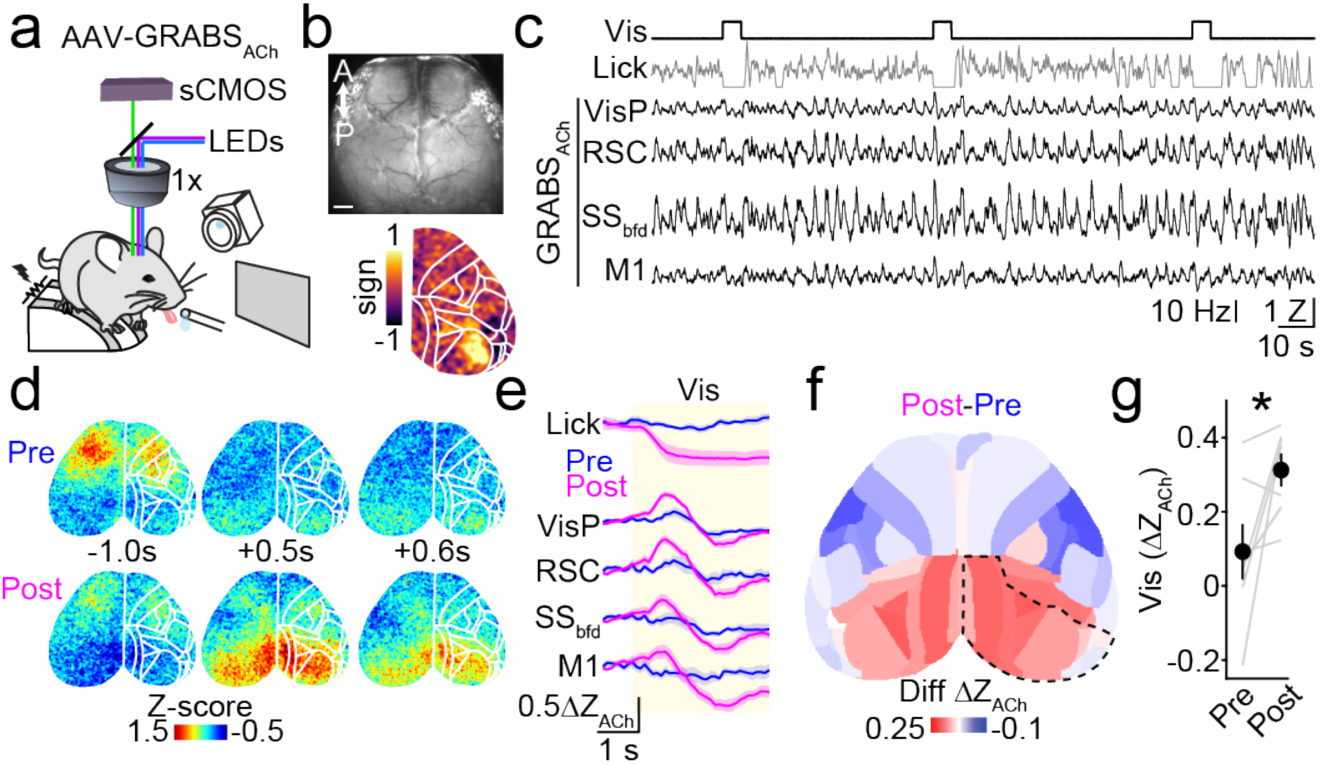
Cholinergic plasticity following visual fear conditioning. (**a**) Schematic of the widefield imaging setup. (**b**) Upper: example transcranial imaging field of view showing GRABS_ACh_ expression. Scale bar: 1mm. Lower: averaged retinotopic field sign maps with Allen CCFv3 parcels overlaid in white. (**c**) Example time series from a portion of a post-conditioning session showing onsets of visual stimuli, lick rate, and traces corresponding to GRABS_ACh_ activity in Allen parcels (VisP; primary visual cortex, RSC; retrosplenial cortex, SS_bfd_; somatosensory cortex barrel field, M1; primary motor cortex). (**d**) Example image frames showing a visual stimulus response pre-(Upper) and post-(Lower) conditioning from the same animal. White lines indicate Allen CCFv3-derived parcellation. Times in seconds are relative to visual stimulus onset at t=0s. White lines indicate Allen CCFv3-derived parcellation. (**e**) Traces show visual stimulus-averaged activity pre-(blue) and post-(magenta) conditioning from the Allen parcels in (**c**) (n=7 mice). (**f**) Average spatial map of the difference (post minus pre-conditioning) in the evoked visual response amplitude for each Allen parcel (n=7 mice). Red indicates increased response amplitude and blue indicates decreased response amplitude. (**g**) Average response across visual parcels, represented by dashed outline in (**f**), pre- and post-conditioning (n=7 mice) * indicates p<0.05, paired t-test. Data in panels **e**, **f**, & **g** are represented as mean ± SEM.

We then examined whether the increased ACh release was also associated with improved perceptual performance. As above, the averaged evoked response in visual cortex was significantly correlated with lick suppression and learning was associated with a significant improvement in the accuracy of a linear decoder to detect stimulus occurrence based on ACh release in visual cortex (Extended Data Figure 4).

These findings suggest a key role for ACh signaling in the functional reconfiguration of cortical activity associated with learning. We therefore tested whether these cholinergic dynamics were necessary for either the expression of conditioned behavior or the enhancement of the cortical visual representation. First, we bilaterally implanted injection cannulas and expressed GCaMP8s using AAV injection in VisP. This approach again allowed us to monitor cortical activity, including retinotopic identification of VisP borders (Figure 5a). We infused the muscarinic cholinergic antagonist scopolamine bilaterally into VisP both before and after training. As above, conditioning produced robust lick suppression and enhancement of the visually evoked cortical response (Figure 5b-c). Prior to conditioning, scopolamine injection did not alter either licking behavior or the evoked cortical signal (Pre vs Post lick index, −0.14±0.14 vs. 0.02±0.08, p=0.35, paired t-test, n=4 mice, Figure 5b). Scopolamine also did not alter baseline licking or behavioral state measured by pupil dilation (Extended Data Figure 5). However, after training scopolamine significantly disrupted lick suppression (F=56.50, p<0.001, ANOVA; p<0.01 Tukey’s Post-hoc for scopolamine vs. vehicle, n=4 mice) and significantly reduced the magnitude of the cortical visual response (Pre scop/vehicle ratio mean, 0.98±0.23, t(3) = 0.09, p=0.93, Post ratio mean, 0.48±0.10, t(3) = 5.17, p=0.01,R^2^ 0.09, one-sample t-test, Figure 5d). We observed similar results in a separate cohort using systemic scopolamine injection (Extended Data Figure 5). Cholinergic signaling via muscarinic receptors in VisP is thus necessary for the expression of both conditioned behavior and the learning-dependent increase in cortical visual representation.

**Figure 5:**
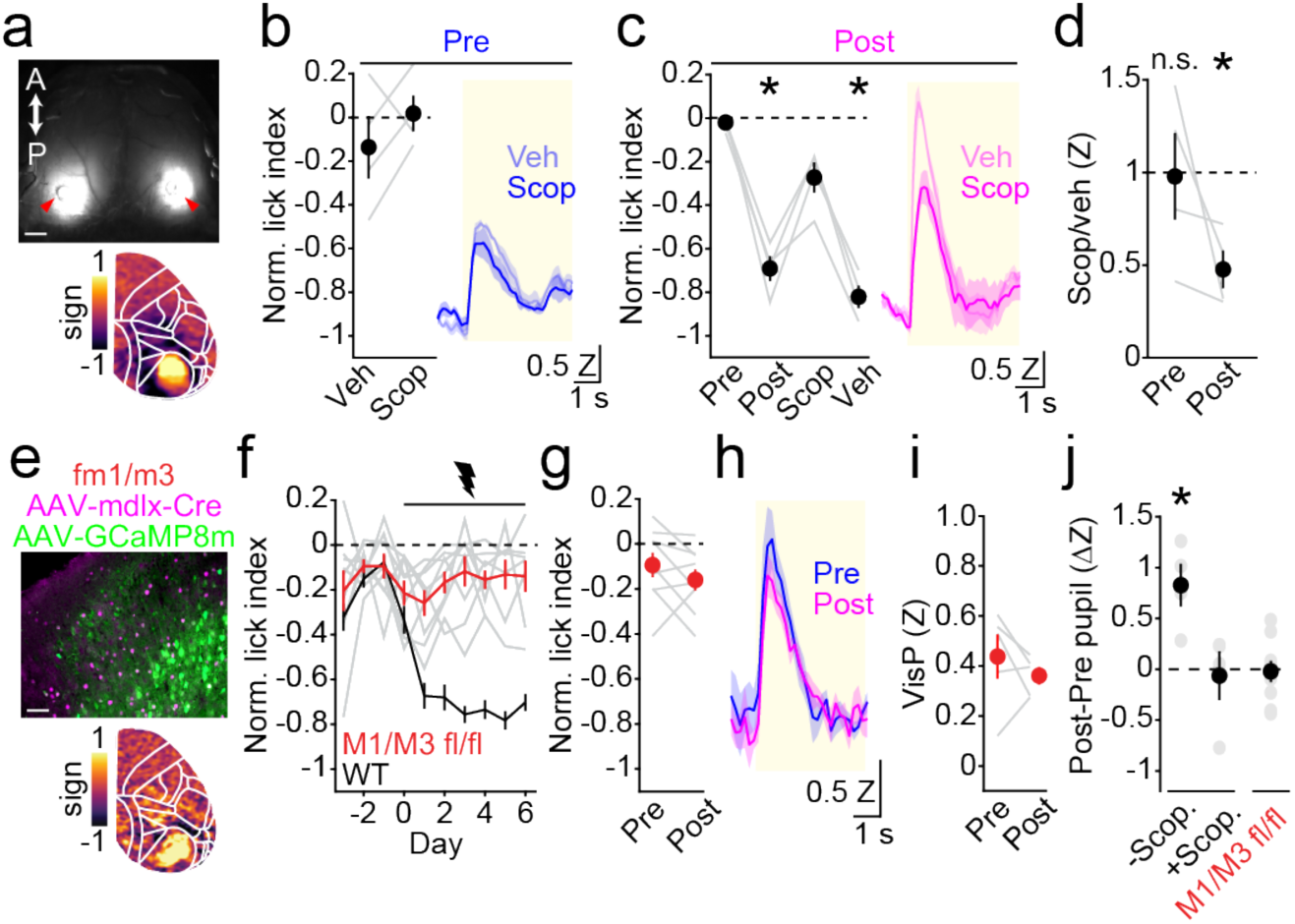
Disrupting cholinergic signaling eliminates conditioned behavior and visually evoked activity. (**a**) Upper: example transcranial imaging field of view showing GCaMP8 expression three weeks after AAV injection. Red arrowheads indicate craniotomy sites for local pharmacology experiments. Scale bar: 1mm. Lower: averaged retinotopic field sign maps with Allen CCFv3 parcels overlaid in white. (**b**) Normalized lick index values before conditioning in the presence of scopolamine (scop, dark blue) or vehicle (light blue). Traces show visual stimulus-averaged activity from the visual cortex before conditioning, after local injection of either scopolamine or vehicle (n=4 mice). (**c**) Average pre-, post-conditioning, scopolamine session, and vehicle session conditioning lick indices. * indicates p<0.05 for pairwise post-hoc comparisons (Tukey-Kramer corrected) against preconditioning session following a significant (p<0.05) main effect of session in repeated measures ANOVA. Traces show visual stimulus-averaged activity from the visual cortex after conditioning following local injections of either scopolamine (dark magenta) or vehicle (light magenta) (n=4 mice). (**d**) Magnitude of visual response (Scopolamine/vehicle) pre- and post-conditioning. * indicates p<0.05 for one-sample t-test comparing population response to hypothesized mean of 1. (**e**) Upper: example confocal image from a M1/M3fl/fl mouse co-injected with mdlx-Cre and camkII-GcaMP8m in visual cortex Cre-expression was visualized using a fluorescent antibody. Lower: averaged retinotopic field sign maps with Allen CCFv3 parcels overlaid in white. (**f**) Normalized lick indices per day, with day 0 as the first session where the conditioned stimulus was paired with foot-shock. Individual M1/M3 fl/fl animals are shown as gray lines (n=9), with the average in red. (**g**) Average pre- and post-conditioning lick indices (n=9 mice). (**h**) Traces show visual stimulus-averaged activity from the visual cortex pre-(blue) and post-(magenta) conditioning (n=5 mice). (i) Average response across the visual cortex pre- and post-conditioning (n=5 mice). (**j**) Post-minus pre-conditioning pupil area difference without scopolamine and in the presence of scopolamine (n=4 mice), and in M1/M3fl/fl mice (n=9 mice). * indicates p<0.05 for one sample t-test against hypothesized mean of zero. Data in panels **b**, **c**, **d**, **f**, **g**, **h**, **i** & **j** are represented as mean ± SEM.

Prior studies suggest that ACh can directly excite L1-INs, providing a mechanism by which enhanced cholinergic release might mediate local circuit disinhibition ^3,79,80^. We therefore asked whether expression of muscarinic receptors in cortical interneurons is necessary for learned behavior and cortical plasticity. We used AAV injection to drive expression of GCaMP8m in cortical PNs and Cre recombinase selectively in GABAergic interneurons of VisP in mice harboring conditional alleles of the Type 1 and Type 3 muscarinic receptors (M1/M3 fl/fl), the primary excitatory receptors in GABAergic cells ^81–83^ (Figure 5e). This approach allowed us to monitor behavior and cortical activity in animals lacking muscarinic excitation of VisP interneurons. In contrast to control animals, deletion of muscarinic signaling abolished the learning of conditioned lick suppression (Pre vs. Post lick index, −0.09±0.05 vs. −0.16±0.05, p=0.22, paired t-test, n=9 mice, Figure 5g). Moreover, imaged animals showed no significant increase in visually evoked cortical activity (Pre vs. Post evoked response, 0.44±0.09 ΔZ vs. 0.36±0.04 ΔZ, p=0.36, paired t-test, n=5 mice, Figure 5h-i). Finally, both intracortical scopolamine injection and deletion of muscarinic receptors from INs in VisP prevented the visually-evoked pupil dilation associated with conditioning (-scop post-minus-pre ΔZ, t(3) = 3.95, p<0.05, +scop post-minus-pre ΔZ, t(3) = 0.26, p=0.81, M1/M3^fl/fl^ post-minus-pre ΔZ, t(8) = 0.20, p=0.81, one-sample t-test, Figure 5j). Overall, these findings also demonstrate that cholinergic signaling in interneurons is necessary for both learning conditioned behavior and the associated plasticity of visual representations in the cortex.

Behaviorally-coupled ACh release is hypothesized to play a critical role in gating learning-related plasticity in cortical circuits. For example, electrical stimulation of the basal forebrain induces shifts in the receptive field structure of neurons in auditory cortex ^2,7,8^. In these studies, ACh was considered to gate subsequent plasticity in cortical circuits, potentially through a temporary relief of GABAergic inhibition that enabled momentary release of firing in auditory cortex ^3^. Moreover, the acquisition of reward timing by visual cortex neurons was shown to require ACh release for the induction of plasticity but was unnecessary for the expression of the learned firing patterns ^14^. In contrast, we observed that visual fear conditioning resulted in sensory-evoked ACh release within visual areas. Muscarinic receptor signaling in GABAergic interneurons within VisP was necessary for both acquisition and expression of conditioned behavior and enhancement of the visual representation. Thus, we find that learning-induced plasticity of ACh release can serve as the primary mechanism of expression for conditioned behavior.

Our observation that visual fear conditioning resulted in a spatially restricted, sensory-induced release of ACh in posterior cortical areas is consistent with our previous work showing that spontaneous fluctuations in behavioral state are associated with dynamic, spatiotemporally heterogeneous activity ^4^. Furthermore, anatomical studies show that distinct populations of cholinergic neurons in the basal forebrain exhibit diverse long-range projection patterns ^84–87^ and functional data demonstrate ACh release in visual areas may exhibit retinotopy ^77^. Our results suggest that learning is associated with plasticity in the cholinergic system, promoting visually-induced ACh release selectively in visual areas and contributing to heterogeneous modulation across the cortex. The mechanisms of this plasticity remain unknown but may reflect circuit coordination between visual and frontal cortex and the basal forebrain and the potential for modification of descending control over cholinergic neuronal activity.

Learning reconfigures the functional organization of cortical GABAergic circuits in both auditory and visual areas ^88–90^, although the mechanisms are largely unexplored. Our results suggest sensory-evoked ACh release as a candidate explanation, in which cholinergic excitation of L1-INs through muscarinic receptor activation ^31,79^ drives inhibition of VIP- and PV-INs and subsequent disinhibition of PN output. L1-INs, and particularly those expressing NDNF, have emerged as a potential state-dependent controller of local cortical circuits ^39,64,65,67,91^ that may be critical for both induction and expression of learning ^3,39,41,67^. We propose that learning-dependent plasticity of ACh release harnesses this stable circuit motif to promote conditioned, visually-guided behavior. Notably, VIP-IN-mediated circuit disinhibition has also been posited as a key motif in cortical circuits ^61–63^. However, our results suggest that, under circumstances of behaviorally-cued ACh release, L1-IN-mediated inhibition of VIP-INs dominates the direct cholinergic excitation of the latter population.

In conclusion, we demonstrate that visually cued fear conditioning results in selective, sensory-evoked ACh release in visual cortex, enhancing sensory representations and promoting visually guided behavior by engaging a disinhibitory GABAergic circuit motif. Thus, plasticity of cholinergic release and dynamic circuit reconfiguration may be a broad mechanism for learning-dependent modification of cortical circuits necessary for behavior.

## Data availability

Data used for all analyses in the present manuscript are provided in the accompanying spreadsheet file.

## Code availability

Code used in analysis is available at GitHub: https://github.com/cardin-higley-lab/Moberly_et_al_2026

## Acknowledgements

The authors thank all members of the Higley and Cardin laboratories for input throughout all stages of this study. We thank Rima Pant and Estela Murillo for generating adeno-associated virus (AAV) vectors. We thank Dr. Bernardo Rudy for the gift of the initial floxed muscarinic receptor mice. This work was supported by funding from the NIH (R01EY022951 and R01EY035127 to J.A.C., R01MH113852 to J.A.C. and M.J.H., R01MH099045, R21MH121841, and DP1EY033975 to M.J.H., F32EY031133, and K99EY035344 to A.H.M, and P30EY026878 to the Yale Vision Core.

## Author contributions

A.H.M, J.A.C., and M.J.H. conceptualized and designed the study. A.H.M. collected and analyzed the data. A.H.M, J.A.C., and M.J.H. wrote the manuscript.

## Corresponding author

Correspondence to Jessica A. Cardin or Michael J. Higley

## Methods

### Animals

Male and female mice (C57BL/6J, Slc17a7-Cre; Jax stock no. 037512 crossed to Ai148; Jax stock no 030238, VIP-IRES-Cre^+/0^; Jax stock no. 031628, SST-IRES-Cre^+/0^; Jax stock no. 018973, PV-IRES-Cre^+/0^; Jax stock no. 008069, M1/M3 fl/fl mice) were kept on a 12h light/dark cycle and provided with ad libitum food and water prior to surgery and/or conditioning experiments. M1/M3 fl/fl mice were produced by crossing floxed M1 muscarinic receptor mice^1^ with floxed M3 receptor mice^2^ to produce double homozygous animals. Animals were individually housed following head-post implants. All behavioral and imaging experiments were performed during the light phase of the cycle. All animal handling and experiments were performed according to the ethical guidelines of the Institutional Animal Care and Use Committee of the Yale University School of Medicine.

### Neonatal sinus injections

In a subset of experimental animals, brain-wide expression of the GRABS_ACh_ Sensor and/or calcium indicators (jGCaMP6s) was achieved via postanal sinus injections as previously described^3,4^. Neonatal pups (postnatal day 0-1) were injected with 8µl (4 µl per hemisphere) with AAV9-Syn-GRABS_ACh_ and/or AAV9-Syn-GCaMP6s-WPRE-SV40 (Addgene no. 100843).

### Surgical procedures

All other surgical procedures were performed on adult mice (between P50 and P180). Mice were anesthetized using 1-2% isoflurane and maintained at 37°C for the duration of the surgery. For widefield imaging implants, the skin and fascia over the skull were removed from the nasal bone to the posterior of the intraparietal bone and laterally between the temporal muscles. The surface of the skull was cleaned with saline and gently polished with wool buffing pads. The edges of the incision were secured to the skull with tissue glue (Vetbond, 3M). A custom titanium headpost was secured to the posterior of the nasal bone with transparent dental cement (C&B Metabond, Butler Schein). A thin layer of cyanoacrylate (Maxi-Cure, Bob Smith Industries) was used to cover the skull and left to cure for at least 30 minutes at room temperature to provide a smooth surface for transcranial imaging.

In IRES-Cre mice, injections of AAV9-Syn-FLEX-GCaMP6s-WPRE-SV40 (Addgene no. 100845, diluted to a titer of 2.5×10^12 vg/mL) were targeted to VisP. For layer 1 imaging experiments, 500 nL of AAV9-mDlx-GCaMP6f-Fishell-2 (Addgene no. 83899, diluted to a titer of 2.2×10^12 vg/mL) were targeted to VisP. A subset of mice was locally injected with AAV9-CamKIIa-jGCaMP8m-WPRE (Addgene no. 176751, diluted to a titer of 2×10^12 vg/mL). In conditional M1/M3 fl/fl mutant mice, CN1851-rAAV-hI56i-minBglobin-iCre-4X2C-WPRE3-BGHpA (500 nL Addgene no. 164450, diluted to a titer of 1.7X10^12 vg/mL) was locally injected into VisP. A subset of these animals (5 out of 9) were co-injected with AAV-minBglobin-iCre and AAV9-CamKIIa-jGCaMP8m-WPRE. All viral injections (600-800 nL) were made via a beveled glass micropipette at a rate of 1-2 nL/second into V1 layer 2/3 (between 150-300 µm in depth) using a Nanoject III (Drummond). After injections, pipettes were left in the brain for more than 10 minutes to prevent backflow.

For two-photon calcium imaging implants, mice were anesthetized and the scalp was cleaned with Betadine and 70% ethanol. The scalp was removed, and the fascia over the skull was cleaned with saline. A custom titanium headpost was secured to the skill with opaque dental cement (C&B Metabond, Butler Schein). A 3 mm^2^ craniotomy was made over V1, and a glass window, consisting of a 3 mm^2^ inner coverslip adhered with ultraviolet-curing adhesive (Norland Optical Adhesives #81) to a 5 mm^2^ outer coverslip (both #1 Warner Instruments), was inserted into the craniotomy and secured with cyanoacrylate glue (Loctite). Metabond was applied to cover any remaining exposed skull. Analgesics were given immediately after surgery (5 mg/kg Carprofen) and on the following two days to aid recovery. Mice were maintained on an antibiotic diet (Sulfatrim, Butler Schein) to prevent infection and allowed to recover for at least 5 days before subsequent handling and behavioral experiments.

### Histology

Histology was performed on a subset of animals at the conclusion of imaging and behavioral experiments to confirm indicator expression. Mice were deeply anesthetized with isoflurane and perfused intracardially with phosphate-buffered saline (PBS) followed by 4% paraformaldehyde in PBS. Brains were postfixed overnight at 4°C and subsequently stored in PBS. Tissue was sectioned at 50 µm using a vibrating blade vibratome (Leica VT 1000 S), mounted onto slides, and visualized via light microscopy. To validate expression of Cre in a subset of M1/M3 fl/fl animals, coronal slices were pretreated in blocking solution (2 percent normal goat serum, 0.1 percent Triton X-100 in PBS) for four hours then incubated with primary antibodies at 1:1000 (rabbit anti-Cre, Sigma 69050) at 4 degrees for 24 hours. The following day, slices were washed with PBS and incubated in secondary antibodies (anti-rabbit-Alexa Fluor 647, Invitrogen) at 1:1000 for two hours at room temperature. Slices were washed 3 times in PBS then mounted on glass slides. Widefield and confocal epifluorescence images and confocal images were taken with a Zeiss LSM 900.

### Widefield imaging

Imaging was performed using a Zeiss Axio Zoom microscope with a 1x, 0.25 numerical aperture objective. Widefield excitation light was from an LED bank (Spectra X Light Engine, Lumencor). Emitted light passed through an image splitter (TwinCam, Cairn Research), then through a 525/50nm (GRABS_Ach_ or GCaMP6s) filter (Chroma) before it reached a sCMOS camera (Orca-Flash V3, Hamamatsu). Images were acquired with an 18-20x digital zoom at 512×512 resolution after 4x pixel binning. Each channel was acquired at 10 Hz with a 35-ms exposure using HCImage software (Hamamatsu). For experiments requiring intrinsic signal imaging of visual responses, LEDs (Thorlabs M530L4) provided backscatter illumination, centered and narrowly filtered at 530 nm, coupled to a 1000 um diameter fiber that terminated 45 degrees incident to the brain surface. Image frames capturing backscatter at 530 nm were acquired at 10 Hz.

### Retinotopic mapping and visual stimulation

Before the start of behavioral experiments, cortical visual areas were mapped using a filtered noise stimulus that drifted across the visual field of head-fixed animals ^5^. All visual stimuli were generated using Psychtoolbox-3 in MATLAB and presented on a gamma-calibrated LCD monitor (Acer V206HQL, 17 inches, spatial resolution of 1600×900, frame refresh rate of 60 Hz, mean luminance of 20 cd/m^2^) positioned 20 cm away from and parallel to the left eye. Filtered noise had a temporal frequency of 2 Hz and a spatial frequency of 0.04 cycles per degree and drifted 20 times along each cardinal axis at rates of 0.086 Hz (altitude) and 0.063 Hz (azimuth). Corrective distortion was applied to the drifting stimuli to alleviate the distortion caused by using a flat monitor to cover a large visual angle. To ensure that stimulus-evoked activity had subsided between sweeps, a gap of 2 seconds was inserted between stimuli. To generate visual field sign maps, we created stimulus-triggered mean ΔF/F_0_ movies using the 2-second inter-sweep interval as the F_0_. A fast Fourier transform (FFT) was used to generate altitude and azimuth phase maps. Then the visual field sign at each pixel was calculated as the sine of the angle between local gradients derived from the phase maps. In conditioning experiments, we presented 5-second, full-screen noise stimuli with the same parameters described above on the same LCD monitor. In a subset of post-conditioning experiments, the contrast of the noise stimulus was randomly varied between one, two, five, ten, and one hundred percent contrast. In discriminative conditioning experiments, the conditioned stimuli were 80-degree drifting gratings (2 Hz, 0.04 cycles per degree) with the CS-plus oriented vertically and the CS-minus oriented horizontally.

### Two-photon imaging

Two-photon imaging was performed using a MOM microscope (Sutter Instruments) coupled to a 16x, 0.8 numerical aperture objective (Nikon). Excitation light was delivered by a titanium-sapphire laser (Mai Tai eHP DeepSee, Spectra-Physics) tuned to 920 nm. Emitted light was collected through a 525/50 nm filter, then a gallium arsenide phosphide photomultiplier tube (Hamamatsu). Images were acquired at 512×512 resolution (with 2x digital zoom) at 30 Hz using a galvo-resonant scan system controlled by ScanImage software (Vidrio). Imaging was performed at a depth of 120-250 µm for layer 2/3 neurons or 20-100 µm for layer 1 neurons. For each animal, only one field of view (FOV) was imaged, chosen to include V1 based on widefield area border maps and blood vessel markers that were observed and registered between functional maps and 2-photon fields of view. Imaging FOVs were matched across days using the template image and z-coordinates from the initial imaging session.

### Behavioral monitoring

All experiments were performed in awake, behaving, head-fixed mice. During widefield and two-photon imaging experiments, the animal’s face was illuminated with an infrared LED bank and imaged with a miniature CMOS camera (Blackfly S USB3, FLIR) with a frame rate of 10 Hz. Video was acquired with SpinView software (FLIR).

### Behavioral conditioning

For all experiments, animals were habituated to being head-fixed for at least 5 days prior to the start of conditioning. During this period, mice were water-restricted to no less than 85% of their initial body weight and trained to lick for rewards (a 1% sucrose solution) at a waterspout coupled to a fiber-optic sensor that detected beam breaks (OPTEX NF-DT05). Licks events were monitored by custom MATLAB software. Detected licks triggered a solenoid valve to deliver a 2µl reward and start a subsequent random time interval (1 to 3 seconds) in which no further reward could be delivered. After mice demonstrated consistent licking throughout head-fixation sessions, we began pre-conditioning sessions in which 10-20 visual stimuli (5-second filtered noise, see visual stimulation section for details) were delivered with an inter-stimulus interval drawn from an exponential distribution (ranging from 50 to 150 seconds, such that mice could not predict the timing of the subsequent trials). After three pre-conditioning sessions, visual stimuli were paired with a 0.5-second, 0.2-0.4 mA foot-shock passed between two grounding braids (Molex) under the animal via a precision-regulated shocker (H13015, Coulbourn Instruments) in two-pole (bipolar) current reversal square wave output mode. The offset of the visual stimulus was followed by a 1-second trace before the onset of the foot-shock. The timing of the shock and visual stimulus was coordinated with custom MATLAB codes through a NI-DAQmx board (PCIe-6315, National Instruments), and all output timing signals were acquired with a Power 1401 digitizer (Cambridge Electronic Design) and sampled at 5 kHz. During conditioning, animals were constantly monitored, and sessions were terminated if signs of stress were observed (e.g., vocalizations, excessive struggling). Mice were trained daily, with sessions lasting approximately 20 minutes.

### Pharmacology

Animals were injected with either sterile saline or scopolamine hydrobromide (1 mg/kg, i.p., Tocris Bioscience; CAS no. 114-49-8) 15 minutes before behavioral sessions. For acute, local injections of muscimol (1 µg/µL, Tocris Bioscience; CAS no. 2763-96-4, dissolved in ACSF), craniotomies were made bilaterally over VisP and covered with silicone sealant (Kwik-cast, WPI) at least one day prior to behavioral sessions. 15 min prior to behavior sessions, mice were head-fixed and injected locally. For acute local scopolamine experiments, craniotomies were targeted to VisP after functional area mapping but prior to behavioral training. During acute injection sessions, 500-600 uL of scopolamine (0.2 mM) was injected into visual cortex at a rate of 1-2 nL/second in awake, head fixed animals, free to run on a wheel under a stereotaxic arm. Animals were left to recover for 15 min in their home cage before head fixation in the behavioral/imaging apparatus.

### Data analysis

All analyses were conducted using custom-written scripts or publically available software packages in MATLAB (Mathworks). No statistical methods were used to predetermine sample sizes. All animals were randomly selected from our colony for inclusion in the study.

#### Preprocessing of behavior data

Pupil videos were cropped from the full-face videos and manually aligned and registered to a common session. Videos were only included if the eye and pupil were clear and unobstructed by the eyelid or surrounding fur. Pupil area was extracted from the cropped videos using FaceMap^6^. Non-normalized pupil area (in image pixels) was used to compare baseline pupil areas. To evaluate visually evoked responses, pupil traces were Z-scored, and the evoked delta calculated by subtracting the mean value from the baseline period.

#### Preprocessing of licking behavior

To measure lick suppression, we used a normalized lick index ^7,8^ comparing the number of licks during the 5-second baseline period preceding the 5-second visual stimulus and the number of licks during the visual stimulus (licks_vis_-licks_baseline_/licks_vis_+licks_baseline_). To estimate lick rates, lick times were convolved with a Gaussian filter (with a 1-second smoothing window), and the lick density was interpolated to match the sampling rate of other simultaneously acquired timing signals (5 kHz).

#### Preprocessing of widefield imaging data

Widefield images were processed as previously described ^3,4^. Briefly, detrending was applied using a low-pass filter (N = 100, fcutoff = 0.001 Hz). Time traces were obtained using (ΔF/F)_i_ = (F_i_ − F_i,0_)/F_i,0_, where F_i_ is the fluorescence of pixel i and F_i,0_ is the corresponding low-pass-filtered signal. To register widefield movies to the Allen Common Coordinate Framework (CCFv3) ^9^, we used field sign maps from visual area mapping sessions to ensure that V1 was well circumscribed by the ‘VisP’ Allen parcel. Subsequent imaging sessions were aligned to the retinotopic mapping session using a rigid, intensity-based registration (imregtform in MATLAB), which utilized raw images from both the retinotopic mapping session and the behavioral imaging sessions. Hemodynamic correction was applied as described in Lohani, Moberly et al. (2022) using the approximate isosbestic excitation of GCaMP6s or GRABS_ACh_ to estimate activity-independent fluctuations in fluorescence. Final activity traces were obtained by z-scoring the corrected ΔF/F signals per pixel. Pixels within Allen Parcels were averaged to yield a final fluorescence trace per parcel. Visual responses for pre- and post-conditioning experiments were evaluated, per parcel, as the difference between peak response during stimulus presentation and the average activity during the preceding three seconds.

#### Cross-parcel correlations

Time-lagged correlations between every pair of Allen CCF parcel traces were computed by performing cross-correlations with a 0.5-second time lag and extracting the maximum correlation coefficient. Parcel-wise correlation matrices were calculated on time series representing all visual stimulus presentation times (vis stim) and time series with the visual stimulus and shock periods removed (spontaneous). The visual stimulus correlation values were subtracted from spontaneous correlation values.

#### SVM classification

To quantify the accuracy with which visual responses are encoded by visual activity, we trained a binary classifier (linear SVM, fitcsvm) to separate the average activity in VisP during the 5-second visual response and the 5 seconds prior to stimulus onset for trials before and after conditioning. We used 10-fold cross-validation and computed the cross-validation loss (classification error) across all folds. Error rates were reported as average accuracy (1 - loss). From these sessions, we also trained binary classifiers on shuffled VisP time series.

#### Quantification of two-photon signals

Analysis of 2-photon imaging data was performed using custom routines in MATLAB. Motion artifacts were corrected using the NoRMCorre software package ^10^, and regions of interest (ROIs) were selected as previously described^11^. All pixels in each ROI were averaged as a measure of fluorescence, and the neuropil signal was subtracted. Visually evoked responses were Z-scored based on the mean and standard deviation of the pre-visual stimulus baseline activity (3 seconds) such that ZF = (F-Baseline mean)/(Baseline standard deviation). Evoked response amplitudes were quantified as the mean over the 5-second visual stimulus. We used linear mixed-effects models for 2P imaging data, due to its nested structure with multiple cells recorded from each subject. We treated experiment type (pre- or post-conditioning) as a fixed effect, and individuals (mice) as a random effect. We used MATLAB’s fitlme function with the formula: evoked response ∼ 1 + Condition + (1 | mouse: subjectID).

## Extended Data Figures

**Extended Data Figure 1:**
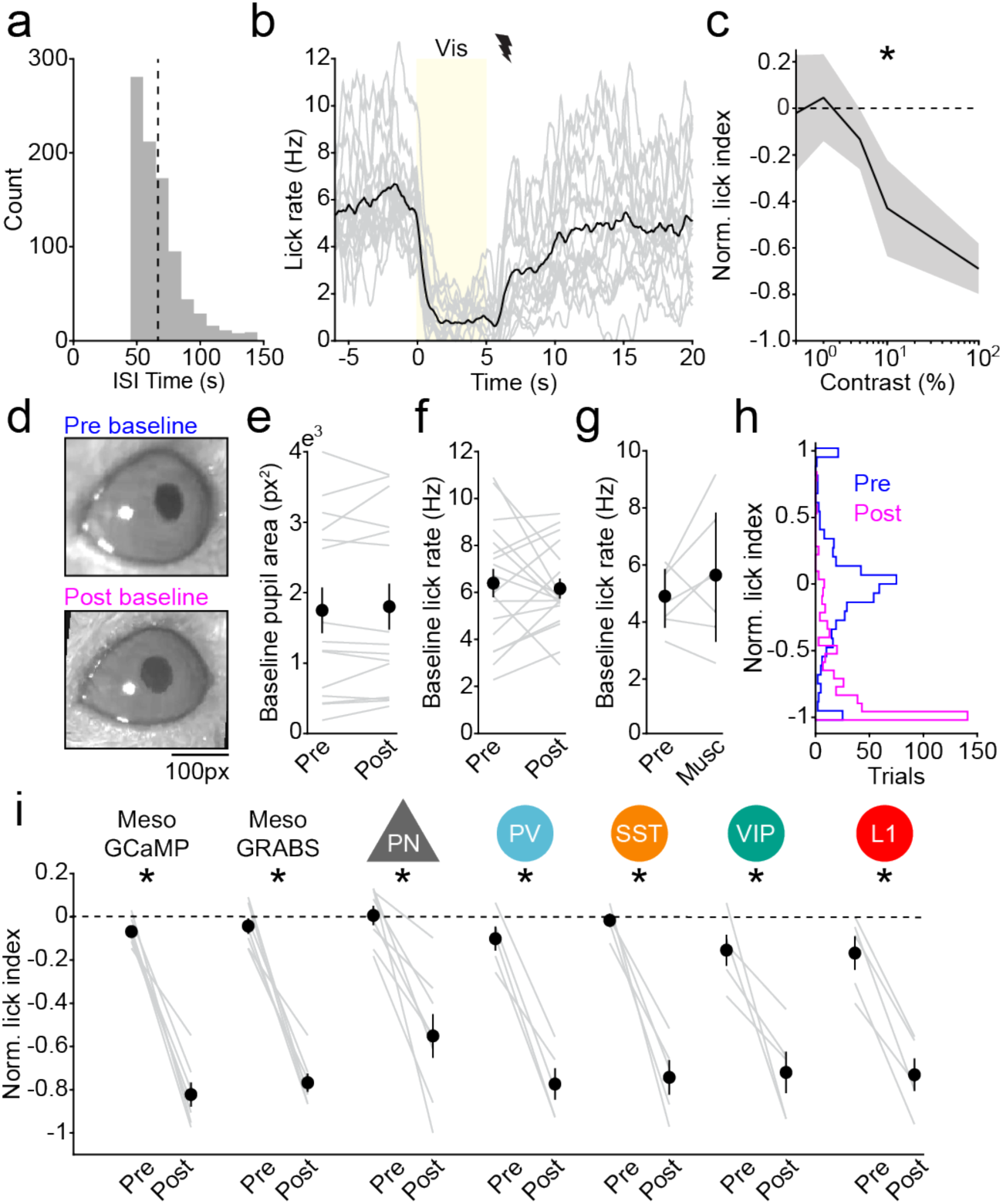
Baseline arousal metrics and behavior summary. (**a**) Histogram of interstimulus intervals across pre- and post-conditioning sessions; dashed line indicates the mean (55 seconds). (**b**) Average visual-stimulus-aligned lick rate from post-conditioning sessions, showing the timescale of licking recovery following foot-shock. Each line represents the average from an individual mouse (n=17 mice). (**c**) Normalized lick index as a function of stimulus contrast (n=3 mice). * indicates a significant main effect of contrast in a mixed-effects model. (**d**) Example pupil images (averaged across the 5-second baseline period) from a pre-(Upper) and post-(Lower) conditioning session. (**e**) Pupil area from the baseline period in pre- and post-conditioning sessions (n=15 mice). (**f**) Baseline lick rates (average from the 5 seconds before visual stimulus onset) in pre- and post-conditioning sessions (n=17 mice). (**g**) Baseline lick rates during pre-conditioning and muscimol treatment sessions (n=7 mice). (**h**) Distribution of normalized lick index values across pre-(blue) and post-(magenta) conditioning sessions (534 and 428 trials, respectively) used in imaging experiments. (**i**) Normalized lick indices pre- and post-conditioning for mice used in 1-photon (meso) and 2-photon imaging experiments. * indicates p<0.05, paired t-test. Data in panels **c**, **e**, **f**, **g** & **i** are represented as mean ± SEM.

**Extended Data Figure 2:**
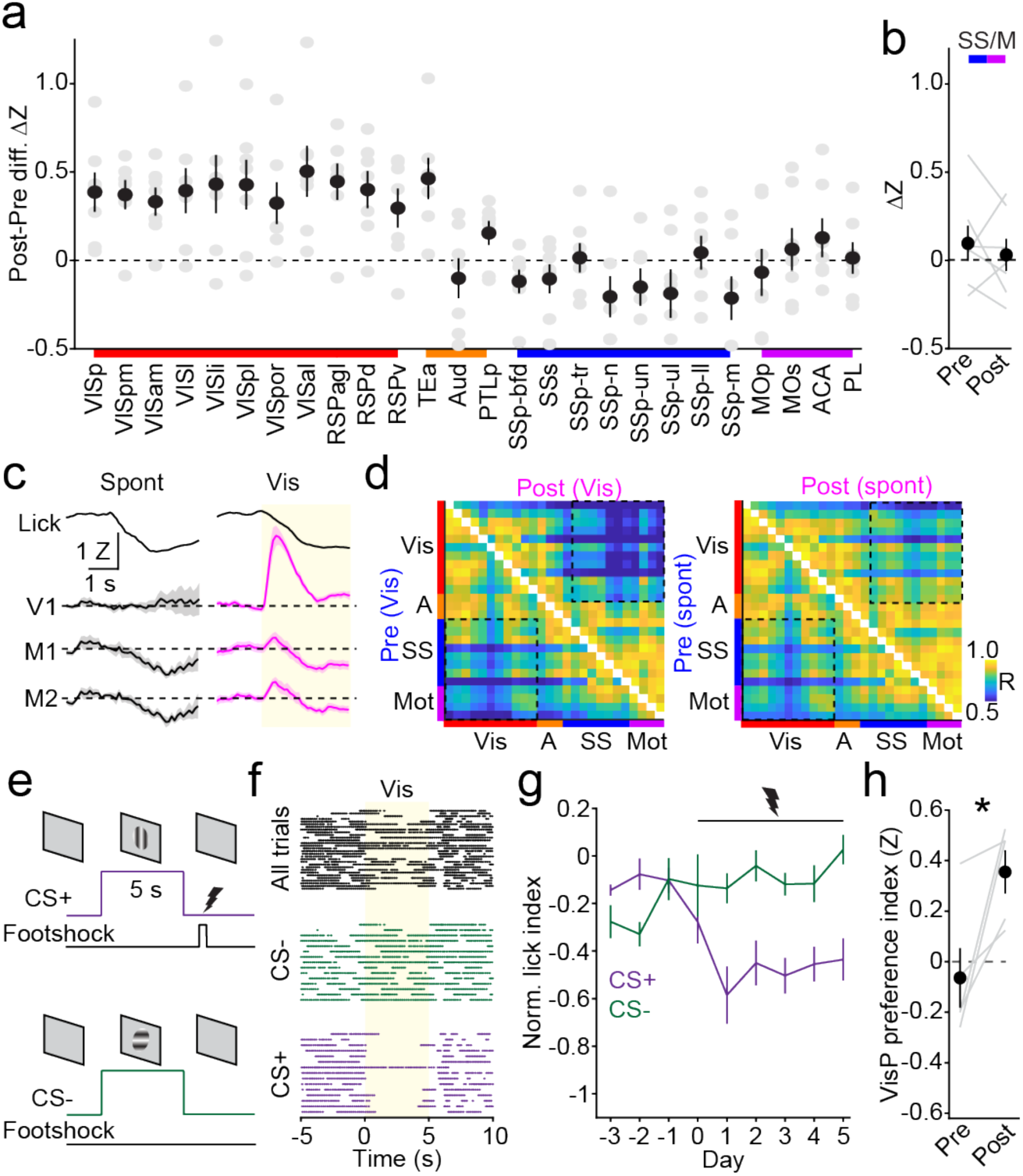
Stimulus-specific plasticity in visual responses. (**a**) Difference (post minus pre-conditioning) in visual response amplitude for all Allen parcels. (**b**) Average visual response pre- vs. post-conditioning for motor and somatosensory parcels. (**c**) Mean traces from VisP and motor (M1 and M2) parcels aligned to spontaneous lick offsets during lick-training compared to post-conditioning sessions with visually-evoked lick suppression (n=7 mice). (**d**) Left: average pairwise correlation matrices during visual stimuli pre- and post-conditioning. Right: average pairwise correlation matrices during interstimulus intervals pre- and post-conditioning (n=7 mice). (**e**) Schematic for differentially-cued fear conditioning experiments. (**f**) Top: example lick raster aligned to visual stimulus onset from a post-conditioning session. Below, trials are split into CS-minus (middle) or CS-plus (bottom) trials. (**g**) Normalized lick indices per day, with day 0 as the first session where the CS-Plus was paired with footshock (n=9 mice). (**h**) Preference index (CS+-CS− / CS++CS−) for visual response in VisP parcel pre- and post-conditioning. * indicates p<0.05, paired t-test. Data in panels **a**, **b**, **c**, **g** & **h** are represented as mean ± SEM.

**Extended Data Figure 3:**
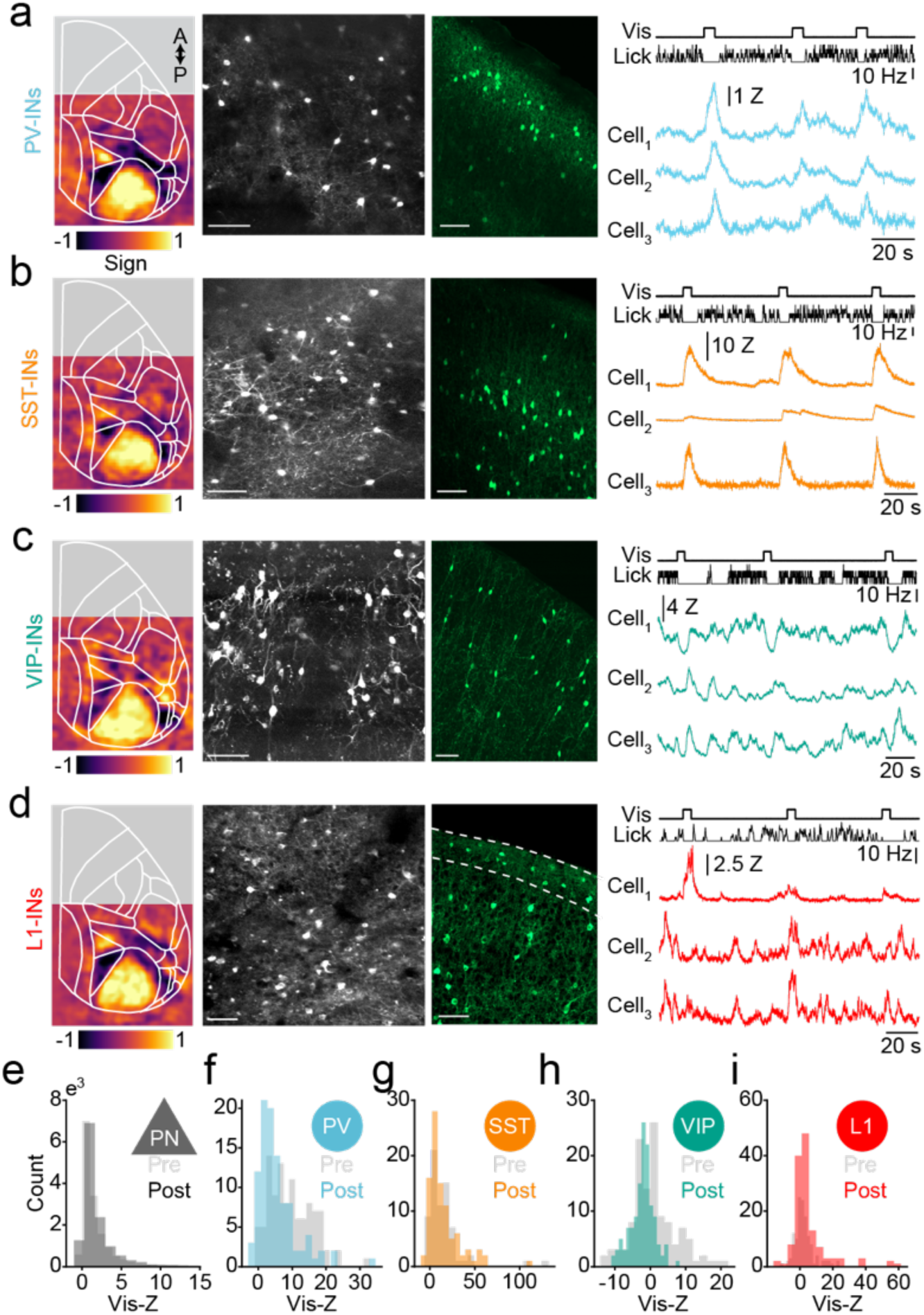
Excitatory and GABAergic imaging summary. (**a**) Left: averaged retinotopic field sign map from GCaMP6s-expressing PV-INs (n=5). Middle two panels: an example 2-photon field of view (scale bar: 50 μm) and a confocal image of GCaMP6s-expressing cells in an example PV-Cre mouse. Scale bar: 70 μm. Right, example time series from one animal from a segment of a post-conditioning session showing onsets of visual stimuli, lick rate, and traces corresponding to selected cell regions of interest. (**b**-**d**) Same as (**a**) for SST-INs, VIP-INs, and L1-INs. Dashed lines indicate layer 1. (**e**) Histograms of average visually evoked cell responses from layer 2/3 PNs (8 mice), PV (5 mice), SST (5 mice), VIP (5 mice), and L1-INs (5 mice).

**Extended Data Figure 4:**
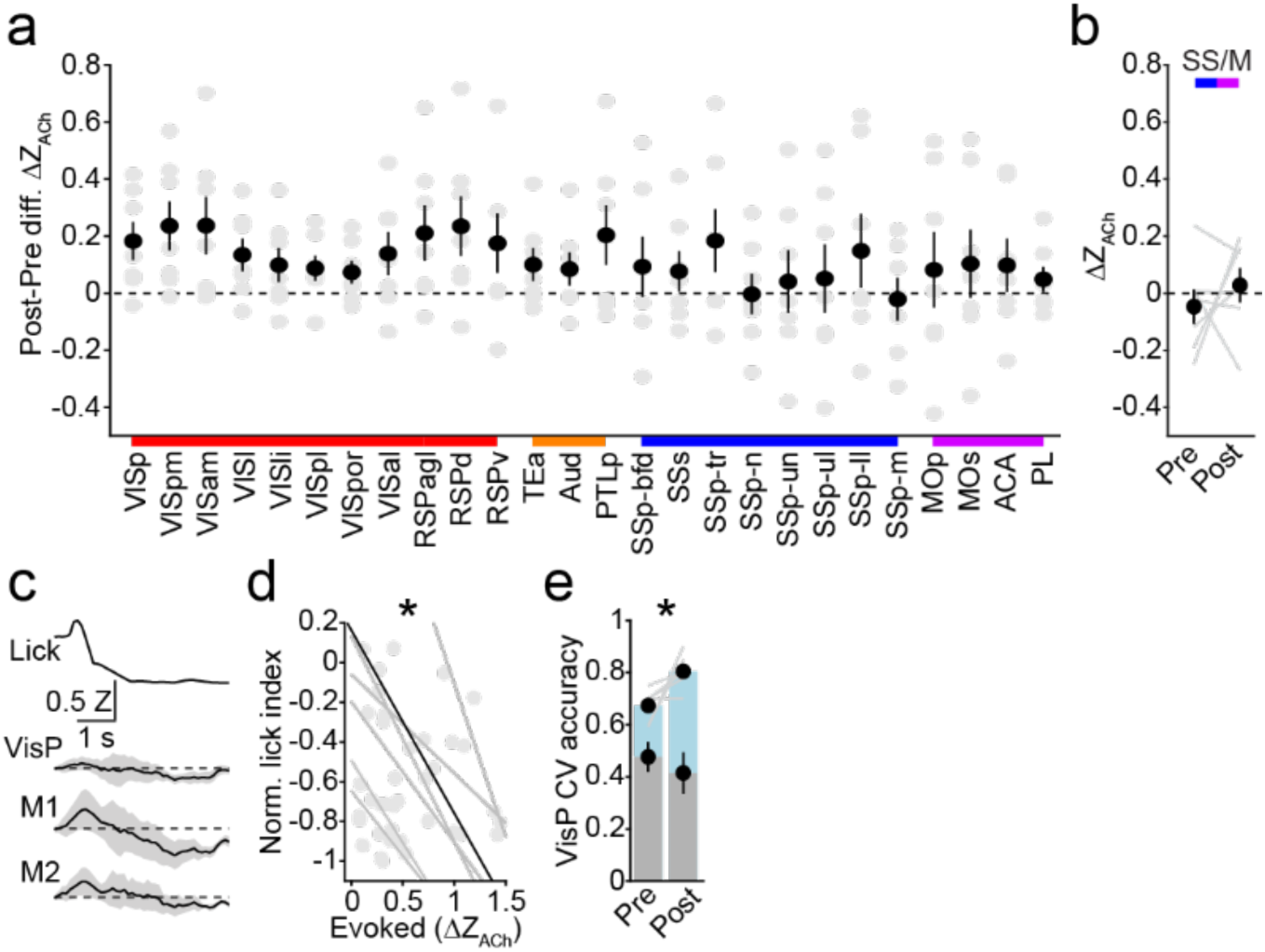
Plasticity in cholinergic signaling. (**a**) Difference (post minus pre-conditioning) in visual response amplitude for all Allen parcels. (**b**) Average visual response pre- vs. post-conditioning in motor (M1 and M2) and somatosensory parcels. (**c**) Mean traces aligned to spontaneous lick offsets in lick-training sessions, showing GRAB_ACh_ response in visual (VisP) and motor (M1 and M2) Allen parcels. (d) Relationship between normalized lick index and evoked visual response; points represent individual sessions with associated best-fit lines. The black line represents the average (n=7 mice). * indicates p<0.05, linear mixed-effects model. (**e**) 10-fold cross-validated accuracy of an SVM trained to classify pre-stimulus baseline activity and visually-evoked activity from the VisP parcel. Gray bars: cross-validated SVM accuracy trained on time-shuffled data. * indicates p<0.05, paired t-test. Data in panels **a**, **b**, **c**, **d**, & **e** are represented as mean ± SEM.

**Extended Data Figure 5:**
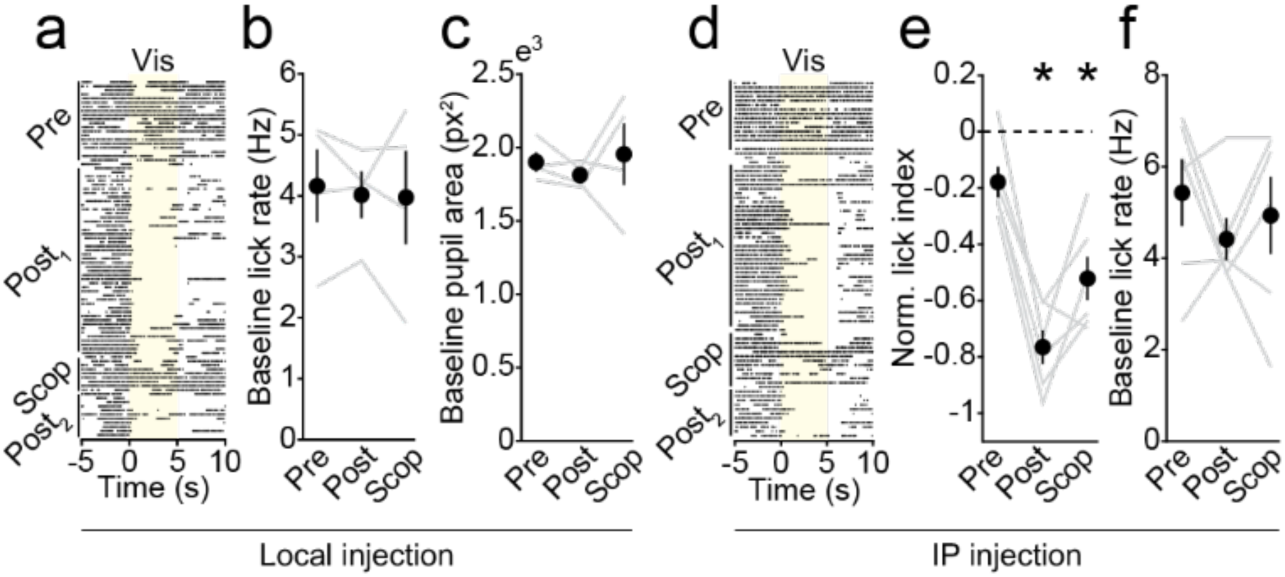
Effects of cholinergic blockers on behavior and baseline activity. (**a**) Example lick raster aligned to visual stimulus onset across pre- and post-conditioning, a scopolamine session (scop), and a post-scopolamine vehicle session. (**b**) Baseline lick rates (averaged over the 5 seconds before visual stimulus onset) in pre-conditioning, post-conditioning, and local scopolamine injection sessions (n=4 mice). (**c**) Pupil area from the baseline period in pre- and post-conditioning sessions and the local scopolamine injection session (n=4 mice). (**d**) Example lick raster aligned to visual stimulus onset across pre- and post-conditioning, an IP scopolamine session, and a vehicle session. (**e**) Average pre- and post-conditioning and IP scopolamine session lick indices (n=6 mice). (**f**) Baseline lick rates (average from the 5 seconds before visual stimulus onset) in pre-conditioning, post-conditioning, and IP scopolamine sessions (n=6 mice). Data in panels **b**, **e**, **c**, & **f** are represented as mean ± SEM.

